# Acentric chromosome congression and alignment on the metaphase plate via kinetochore-independent forces in *Drosophila*

**DOI:** 10.1101/2023.11.14.567057

**Authors:** Hannah Vicars, Travis Karg, Alison Mills, William Sullivan

## Abstract

Chromosome congression and alignment on the metaphase plate involves lateral and microtubule plus-end interactions with the kinetochore. Here we take advantage of our ability to efficiently generate a GFP-marked acentric X chromosome fragment in *Drosophila* neuroblasts to identify forces acting on chromosome arms that drive congression and alignment. We find acentrics efficiently align on the metaphase plate, often more rapidly than kinetochore-bearing chromosomes. Unlike intact chromosomes, the paired sister acentrics oscillate as they move to and reside on the metaphase plate in a plane distinct and significantly further from the main mass of intact chromosomes. Consequently, at anaphase onset acentrics are oriented either parallel or perpendicular to the spindle. Parallel-oriented sisters separate by sliding while those oriented perpendicularly separate via unzipping. This oscillation, together with the fact that in monopolar spindles acentrics are rapidly shunted away from the poles, indicates that distributed plus-end directed forces are primarily responsible for acentric migration. This conclusion is supported by the observation that reduction of EB1 preferentially disrupts acentric alignment. In addition, reduction of Klp3a activity, a gene required for the establishment of pole-to-pole microtubules, preferentially disrupts acentric alignment. Taken together these studies suggest that plus-end forces mediated by the outer pole-to-pole microtubules are primarily responsible for acentric metaphase alignment. Surprisingly, we find that a small fraction of sister acentrics are anti-parallel aligned indicating that the kinetochore is required to ensure parallel alignment of sister chromatids. Finally, we find induction of acentric chromosome fragments results in a global reorganization of the congressed chromosomes into a torus configuration.

**Article Summary:** The kinetochore serves as a site for attaching microtubules and allows for successful alignment, separation, and segregation of replicated sister chromosomes during cell division. However, previous studies have revealed that sister chromosomes without kinetochores (acentrics) often align to the metaphase plate, undergo separation and segregation, and are properly transmitted to daughter cells. In this study, we discuss the forces acting on chromosomes, independent of the kinetochore, underlying their successful alignment in early mitosis.

## Introduction

Chromosome fragments lacking a centromere and a telomere are known as acentrics. Due to their lack of a kinetochore, acentrics are unable to make canonical attachments to microtubules. Kinetochore-microtubule interactions play key roles in mediating chromosome congression to the spindle equator, alignment on the metaphase plate, sister chromosome separation during the metaphase-to-anaphase transition and segregation during anaphase [Navarro & Cheeseman, 2021]. Thus, acentrics might be expected to fail to align properly on the metaphase plate, display separation and segregation defects in anaphase resulting in exclusion from daughter nuclei and the formation of micronuclei [Kanda & Wahl, 2000; LaFountain et al., 2001; Fenech et al., 2011]. However, these expectations are contradicted by the finding that acentric chromosome fragments often display proper sister chromosome separation, poleward migration, and inclusion into daughter nuclei [Royou et al., 2010; Bajer, 1958; Kanda et al., 1998; Vicars et al., 2021; Karg et al., 2017; Bretscher & Fox, 2016; Karg et al., 2015; Warecki & Sullivan, 2018; Khodjakov et al., 1996; Liang et al., 1993; Warecki et al., 2020]. Proposed mechanisms include neo-centromere formation and direct association of acentrics with microtubules or a kinetochore-bearing chromosome [Ishii et al., 2008; Platero et al., 1999, Kanda et al., 2001; Ohno et al., 2016; Karg et al., 2017; Warecki & Sullivan, 2020]. Studies have shown that separation of sister acentric chromosomes during early anaphase relies on Topoisomerase II activity as well as microtubule plus-end pushing forces [Vicars et al., 2021]. Additional work has shown that the anaphase poleward segregation of acentric chromosomes involves Klp3a mediated microtubule-based movement [Karg et al., 2017].

Here we explore the mechanisms that drive congression and alignment of acentric chromosome fragments at the metaphase plate as this also provides insight into the non-kinetochore forces driving congression of intact chromosomes. As with sister chromosome separation and segregation, the kinetochore plays a key role in chromosome congression and alignment on the metaphase plate [Barisic et al., 2014; Ye et al., 2015; Chmatal et al., 2015]. Following nuclear-envelope breakdown, chromosomes positioned within the microtubule arc emanating from opposing centrosomes quickly establish lateral interactions with the plus-end kinetochore-associated motor protein CENP-E (kinesin-7) and are transported to the metaphase plate [Yen, 1992; McEwen et al., 2001; Kapoor et al., 2006]. This is referred to as direct congression. Polar ejection forces mediated by plus-end microtubule dynamics and chromokinesins acting on the chromosomes also drive chromosomes away from the poles toward the equator [Maiato et al., 2017; Tanaka, 2008]. Once at the equator, lateral interactions are converted to plus-end microtubule interactions and the sister kinetochores establish biorientation with microtubules attached to opposing poles [Krenn & Musacchio, 2015]. Opposing forces at the sister kinetochores and chromosome arms result in oscillations but maintain the chromosomes at the metaphase plate. Congression of chromosomes located at the spindle periphery and outside of the arc described by the opposing centrosomes and the metaphase plate require an additional step, known as peripheral congression. Chromosomes must first rely on kinetochore-associated Dynein for microtubule minus-end directed transport to the spindle pole. Once at the pole, the chromosomes engage kinetochore kinesin CENP-E for transport to the metaphase plate. [McEwen et al., 2001; Kapoor et al., 2006].

Despite the key role of the kinetochore in driving chromosome movement, acentric chromosome fragments are capable of moving toward and aligning on the metaphase plate [Carlson, 1938a; Royou et al., 2010; Kaye et al., 2004; Bajer, 1958; Vicars et al., 2021; Karg et al., 2017; Rieder et al., 1986]. This was dramatically demonstrated through live analysis of acentric fragments generated via laser ablation. These fragments rapidly moved away from the poles at a rate similar to intact chromosomes [Rieder et al., 1986]. Subsequent live analysis of X-chromosome acentric fragments generated through endonuclease induction in *Drosophila* neuroblasts revealed they also experience robust anti-poleward forces and are capable of aligning on the metaphase plate [Royou et al., 2010; Vicars et al., 2021; Karg et al., 2017; Warecki et al., 2020].

Left unresolved are the mechanisms driving congression and metaphase plate alignment of chromosome fragments lacking a kinetochore. Here we explore this issue by taking advantage of our ability to efficiently generate GFP-labeled acentric fragments in the genetically tractable *Drosophila* neuroblasts. The GFP tag enables us to distinguish the acentric from the kinetochore bearing chromosomes and facilitates tracking of the acentric as it moves to and aligns on the metaphase plate. We find acentrics efficiently align on the metaphase plate more rapidly than kinetochore-bearing chromosomes. Multiple lines of evidence indicate that proper acentric alignment on the metaphase plate relies on pole-to-pole microtubules and plus-end directed microtubule-based forces. Surprisingly, we find that a small but significant fraction of sister acentrics exhibit an anti-parallel alignment. This suggests that the kinetochore is required to ensure parallel alignment of sister chromatids. In addition, we discovered that induction of acentric chromosome fragments results in a global reorganization of the congressed chromosomes into a torus configuration.

## Results

### Acentric sister chromatids undergo rapid movement at the metaphase plate

As previously described, acentric chromosome fragments have a remarkable ability to congress to the metaphase plate, separate and segregate from one another in anaphase, and incorporate into the reforming daughter nuclei, all while lacking canonical kinetochore-microtubule attachments [Vicars et al., 2021; Warecki et al., 2020]. However, it has been difficult to track an unmarked acentric chromosome fragment during congression and metaphase alignment when the acentric is embedded within the entire chromosome complement. Here we address this issue by taking advantage of a GFP-marker that specifically tags the *Drosophila* X chromosome. In *Drosophila*, the male-specific lethal (MSL3) complex plays a major role in dosage compensation by upregulating genes on the male X chromosome [Sural et al., 2008]. MSL3-GFP preferentially binds the male X euchromatin and thus is suitable for distinguishing the X chromosome from the other chromosomes in live studies. Acentrics are generated using heat-shock-induced expression of the I-CreI endonuclease, which targets rDNA repeats embedded in the X-chromosome centric heterochromatin [Rong et al., 2002; Maggert & Golic, 2005; Paredes & Maggert, 2009; Golic & Golic, 2011]. This produces a kinetochore-bearing heterochromatic chromosome fragment and a euchromatic acentric chromosome fragment. As MSL3-GFP specifically localizes to the euchromatin, it preferentially labels the acentric chromosome fragment and is readily tracked over the course of a mitotic division (Figure 1, Video S1). The MSL3-GFP marker does not disrupt acentric chromosome alignment, segregation, or micronuclei formation when compared to control cells with acentrics alone. In control cells expressing MSL3-GFP but not I-CreI, all of the intact X-chromosomes align normally (N=10). Upon I-CreI induction, 95% (N=18) of the GFP-marked acentrics align on the metaphase plate like intact chromosomes.

**Figure 1:**
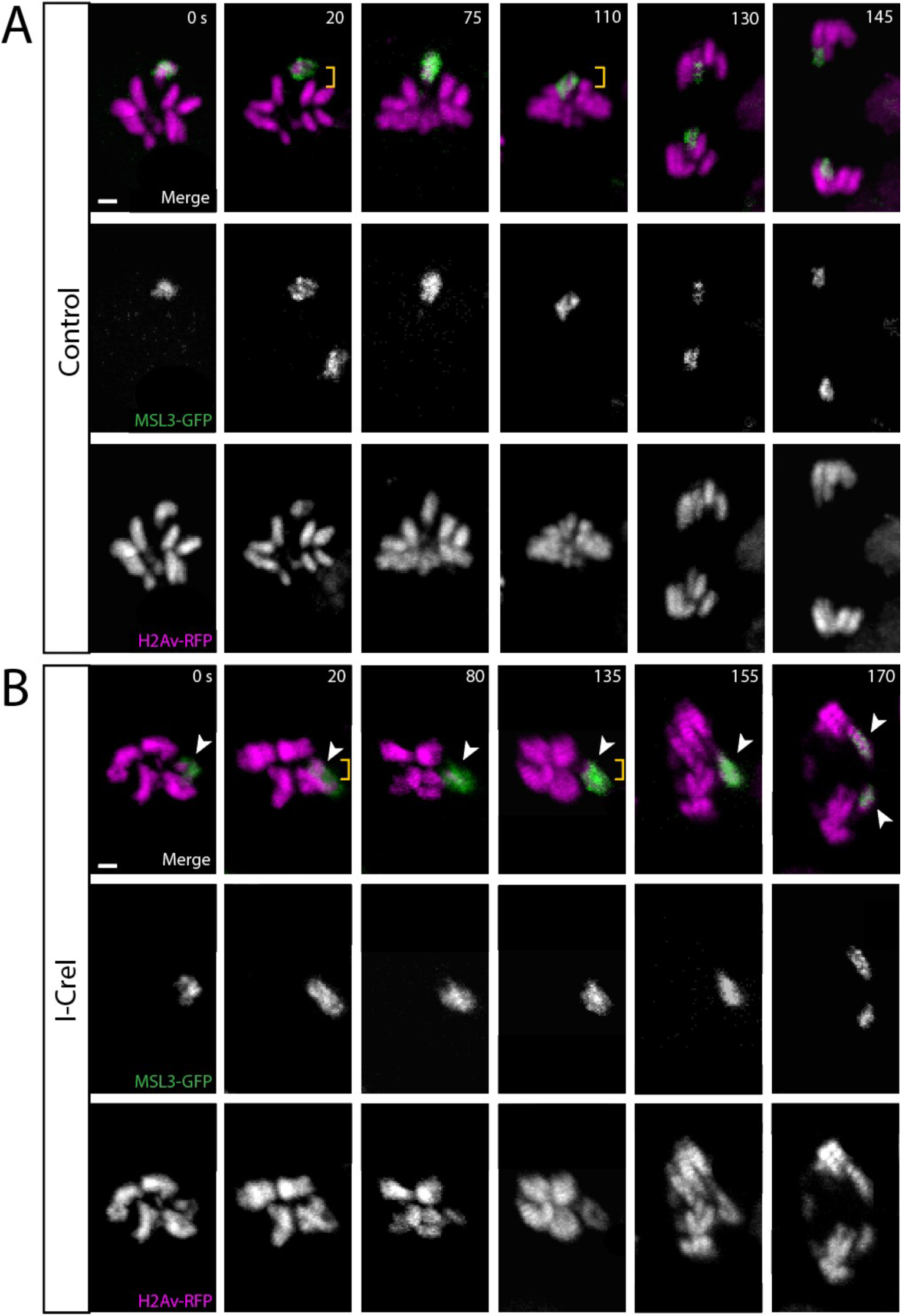
Acentrics align on the metaphase plate. (A) Still frames of a time-lapse movie of a mitotic neuroblast labeled with H2Av-RFP (magenta) and MSL3-GFP (green) not expressing I-CreI. (B) Still frames of a time-lapse movie of a mitotic neuroblast with I-CreI induced acentrics (Video S1). Sister acentrics (white arrowheads) undergo movement at the metaphase plate while aligning (yellow brackets), lag on the spindle equator but eventually separate, segregate, and are incorporated into daughter nuclei. Bars, 2 μm. Time in seconds.

Live analysis 80 seconds prior to anaphase onset was performed to determine the rate on which intact and acentric chromosomes move to the metaphase plate (Table 1, see accompanying schematic). This analysis reveals that on average acentric X chromosomes move at a significantly faster rate while actively aligning at the metaphase plate (11.7 ± 3.5 nm/s, N=14) when compared to intact X chromosomes (8.4 ± 3.3 nm/s, N=12, P=0.01, Mann-Whitney U Test) (Table 1). Intact autosomes (8.5 ± 2.4 nm/s, N=28) align at the metaphase plate at a rate in accordance with the intact X chromosomes (8.4 ± 3.3 nm/s, N=12, P=0.6, Mann-Whitney U Test) (Table 1).

**Table 1:**
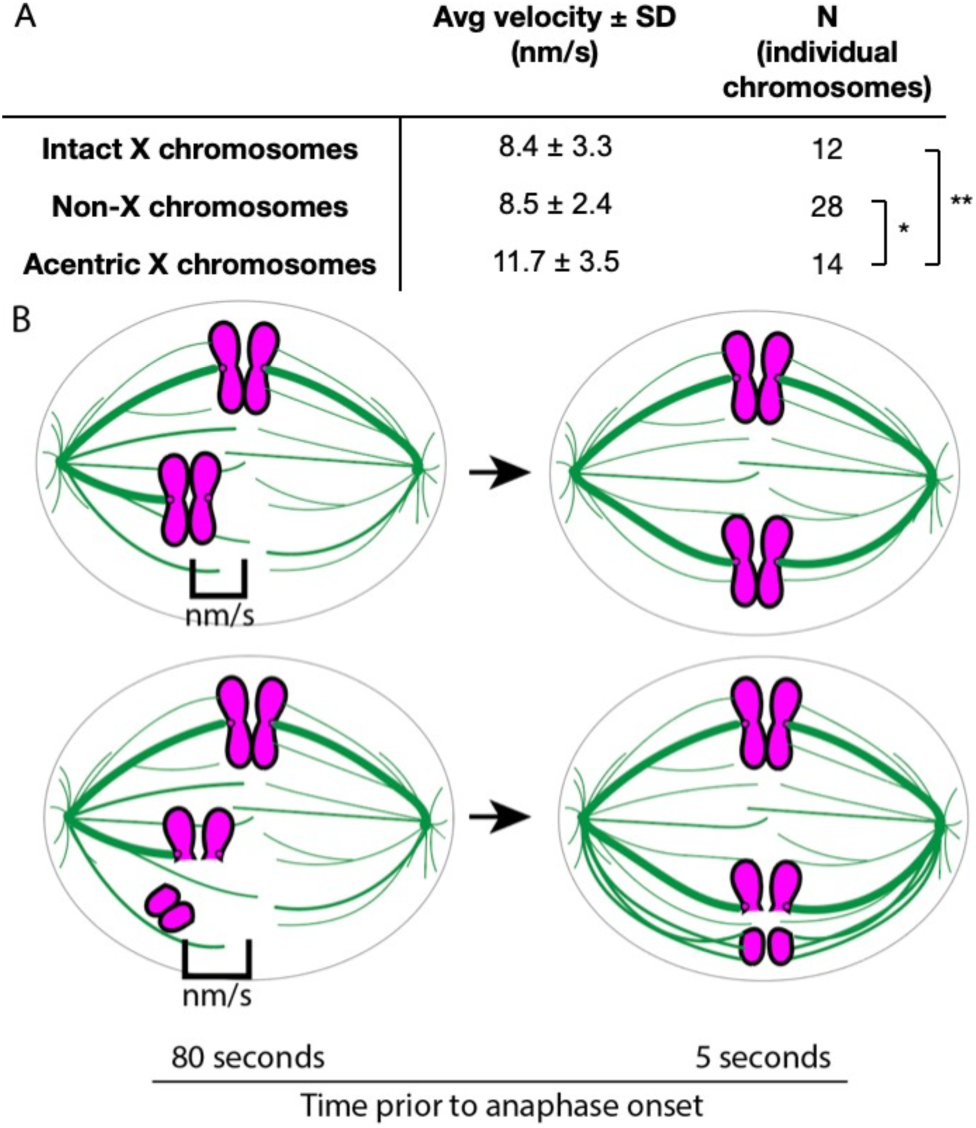
Average velocities of chromosomes at the metaphase plate. (A) Average velocities of individual chromosomes as they align at the metaphase plate. Velocity is measured in nanometers per second. Asterisks indicate statistical significance (*P=0.005, **P=0.01) as determined by a two-sided Mann-Whitney U test. (B) Schematic showing how chromosome velocity is determined. Velocity is defined as beginning 80 seconds prior to anaphase onset and ends one frame (5 s) prior to anaphase onset.

### Acentric induction causes a global reorganization of congressed chromosomes such that acentrics localize to the periphery of the main chromosome mass

Three-dimensional rendering of metaphase images of labeled intact X chromosomes reveal that they are aligned with the autosomes in a planar columnar configuration (Figure 2A, N=5, Video S2). In addition, the intact X chromosome is consistently positioned at the outer edge of the column of congressed chromosomes (Figure 2A, N=5, Video S2). Unexpectedly, we found that I-CreI induction of the acentric X chromosome results in a global reorganization of the congressed chromosomes. Rather than organized in a columnar configuration, the chromosomes are organized into a distinct torus with an absence of chromatin in the center (Figure 2B, N=5, Video S3). We quantified this global organization by measuring the circularity of the entire congressed mass of chromosomes at metaphase. Using the circularity function in ImageJ, we define perfect circles as having a circularity of 1. The circularity measurements demonstrate that the columnar formations in cells in which I-CreI is not expressed are significantly distinct from the torus shapes seen in neuroblasts expressing I-CreI (Figure 2C). The average circularity of metaphase congressed chromosomes with and without acentrics is 0.95 (SD=0.013, N=6) and 0.44 (SD=0.183, N=6), respectively (Figure 2C).

**Figure 2:**
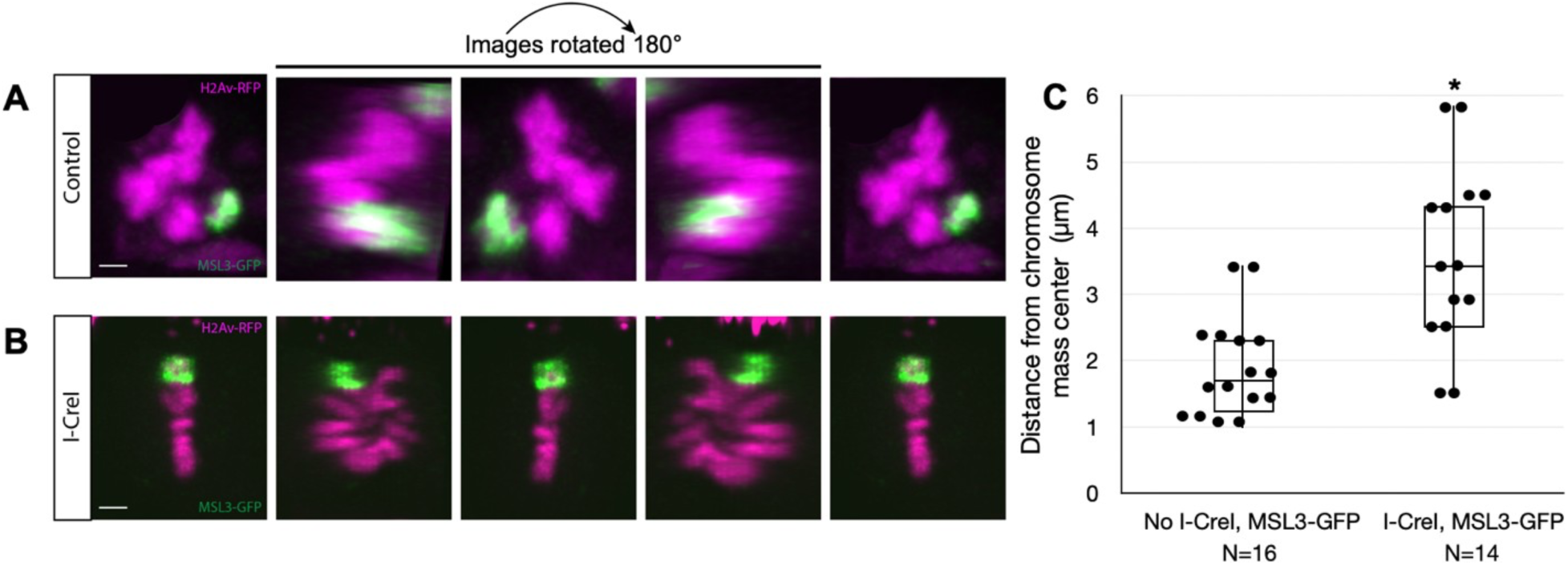
Acentric induction induces a global reorganization of congressed chromosomes such that the acentric localizes to the periphery of the main chromosome mass. (A) Still images of a 3D rendering of a neuroblast at metaphase labeled with H2Av-RFP (magenta) and an intact MSL3-GFP (green) tagged X chromosome (I-CreI not induced) (Video S2). (B) Still frames of a 3D rendering of a neuroblast at metaphase with I-CreI induced acentrics (Video S3). This results in a dramatic global reorganization of the congressed chromosomes into a ring with an absence of chromatin in the center. The acentric localizes to the outer most edge of the chromosome ring. Bars, 2 μm. Images are rotated 180°. (C) Box plot showing the distances of X chromosomes from the chromosome mass center. Distances are measured in µm. Acentric X chromosomes are positioned significantly farther from the chromosome mass center when compared to the intact X chromosome (*P=0.01, Mann-Whitney Test).

As with the intact X chromosome, the acentric is peripherally localized. However, the extent to which the acentric X is positioned away from the main chromosome mass is much greater than that of the intact X chromosome. We quantified peripheral positioning by calculating the distance of the intact X chromosome and acentric from the center of chromosome mass (Figure 2C). On average, acentric X chromosomes are positioned significantly farther (3.60 ± 1.34 µm, N=14) from the chromosomal mass center when compared to intact X chromosomes (1.89 ± 0.74 µm, N=16, P=0.01, Mann-Whitney U Test). The center of the mass of chromosomes is determined by defining the upper, lower, left, and right boundaries of the chromosome mass and calculating the center point. Thus, while the acentric X chromosome fragments efficiently congress to the metaphase plate, they reside in a position distinct from the main chromosome complement (Figure 2).

### Acentrics oscillate while aligning on the metaphase plate

In prometaphase, kinetochore-bearing replicated sister chromosomes remain paired while congressing to the metaphase plate. This is due to the presence of cohesin protein complexes and DNA catenations between sisters [Guacci et al., 1997; Michaelis et al., 1997; Porter & Farr, 2004]. Similarly, sister acentric X chromosome fragments travel to the metaphase plate as a pair, held together via cohesin and DNA catenations [Vicars et al., 2021].

Previous work demonstrated that the orientation of sister acentrics at the metaphase-to-anaphase transition correlates with their mode of separation during late anaphase [Vicars et al., 2021]. Congressed intact chromosomes always align on the metaphase plate perpendicular to the spindle. In contrast, sister acentrics align either perpendicular or parallel to the spindle. Sister acentrics aligned perpendicular to the spindle, separate via unzipping from one another, while those aligned parallel to the spindle slide past one another to opposing poles. Here we provide insight into the trajectories and mechanisms that result in acentrics reaching the plate in multiple orientations. We discovered that acentrics oscillate while aligning on the metaphase plate. To quantify this, we measured the orientation angles of acentric and intact X chromosomes with respect to the cell equator at 5 second intervals during the 80-second interval prior to anaphase onset (Figure 3).

**Figure 3:**
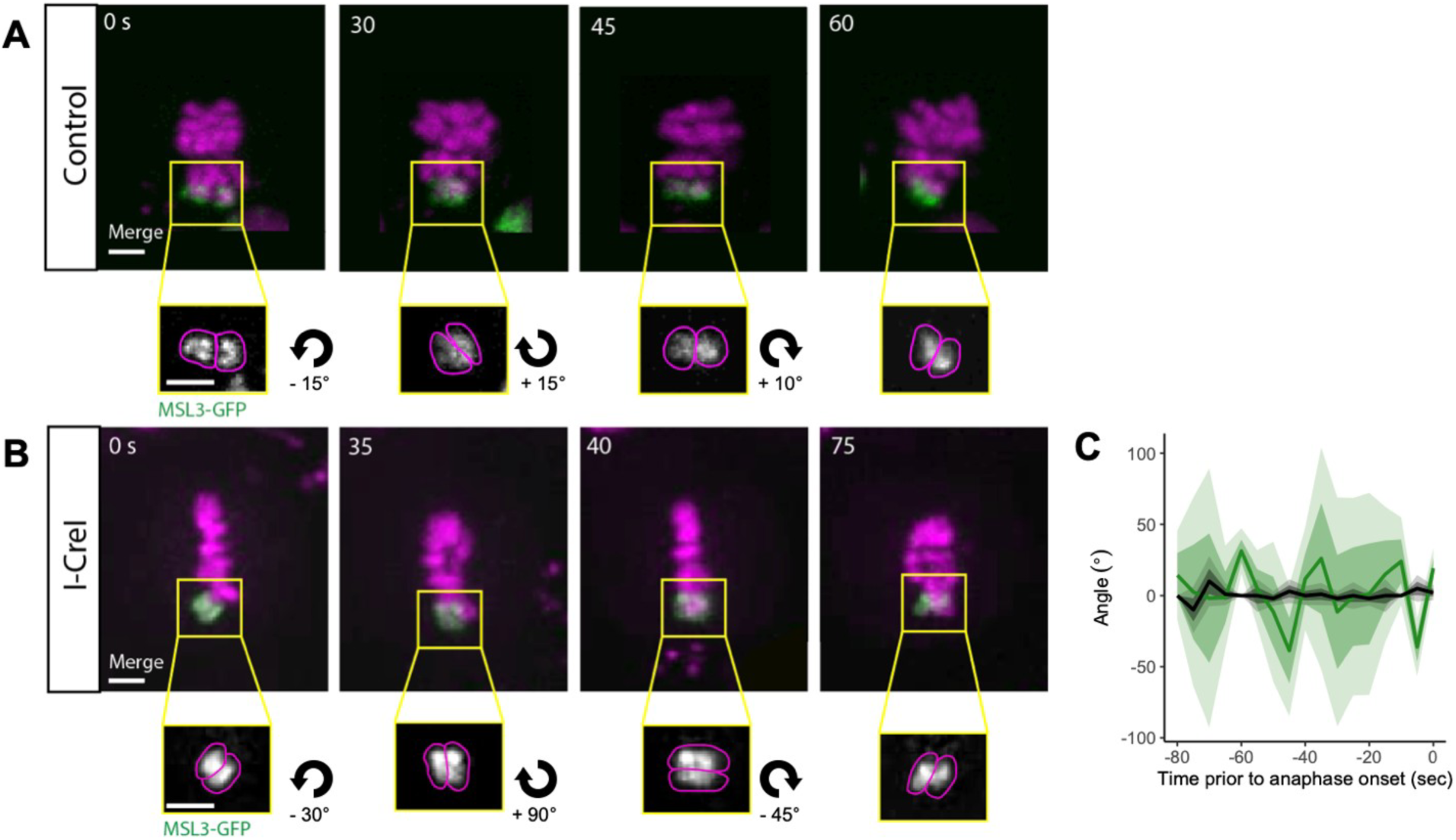
Acentrics undergo varied orientations during metaphase. (A) Still frames of a time-lapse movie of a mitotic neuroblast labeled with H2Av-RFP (magenta) and MSL3-GFP (green), not expressing I-CreI, during metaphase. Zoomed in images show only the X chromosomes marked by MSL3-GFP (gray). (B) Still frames of a time-lapse movie of a mitotic neuroblast with I-CreI induced acentrics during Metaphase (Video S4). Zoomed in images show only the X chromosomes marked by MSL3-GFP (gray). Bars, 2 μm. Time in seconds. (C) Line graph showing the change in orientation of acentric (green) and intact (black) X chromosomes over time, prior to anaphase onset. Data points every 5 seconds. Shaded regions represent one- and two-times the standard deviation.

Kinetochore-bearing sister chromatids align at the metaphase plate undergoing no oscillations, most often remaining perpendicular to the mitotic spindle (Figure 3A, 3C). However, acentric sisters oscillate as they line up at the metaphase plate, alternating between orienting parallel and perpendicular to the spindle and division axis (Figure 3B, 3C, Video S4). These results suggest that the parallel and perpendicular orientations of the acentric at the metaphase plate are due to the fact that they oscillate while aligning on the plate prior to anaphase onset. During anaphase, acentric sisters that are oriented parallel separate by sliding, while those oriented perpendicularly separate via unzipping [Vicars et al., 2021].

### Anti-parallel alignment of sister acentrics

To determine acentric orientation more precisely during metaphase alignment, neuroblasts expressing inducible I-CreI, the histone marker H2Av-RFP, and the telomere marker HOAP-GFP [Cenci et al., 2003] were imaged live. Marked telomeres enabled us to determine the orientation of paired sister acentrics with respect to one another as they aligned on the metaphase plate. A surprising outcome of this analysis is that a low but notable frequency of sister acentrics exhibited anti-parallel alignment. As expected, 100% (25/25) of intact chromosomes have telomeres paired and aligned (Figure 4, Video S5). However, in 12% (4/34) of paired sister acentrics, the telomeres are oriented in opposing directions, with the remaining 88% (30/34) exhibiting normal telomere-to-telomere alignment. This implies that stable pairing of sister chromatids is possible despite the absence of a gene-by-gene alignment. These findings indicate the kinetochore plays a key role in ensuring the fidelity of sister chromosome alignment in *Drosophila*. Additionally, we find acentrics are oriented with their telomeres facing the periphery of the mass of congressed chromosomes (N=10, Figure 4C & 4D, Video S6).

**Figure 4:**
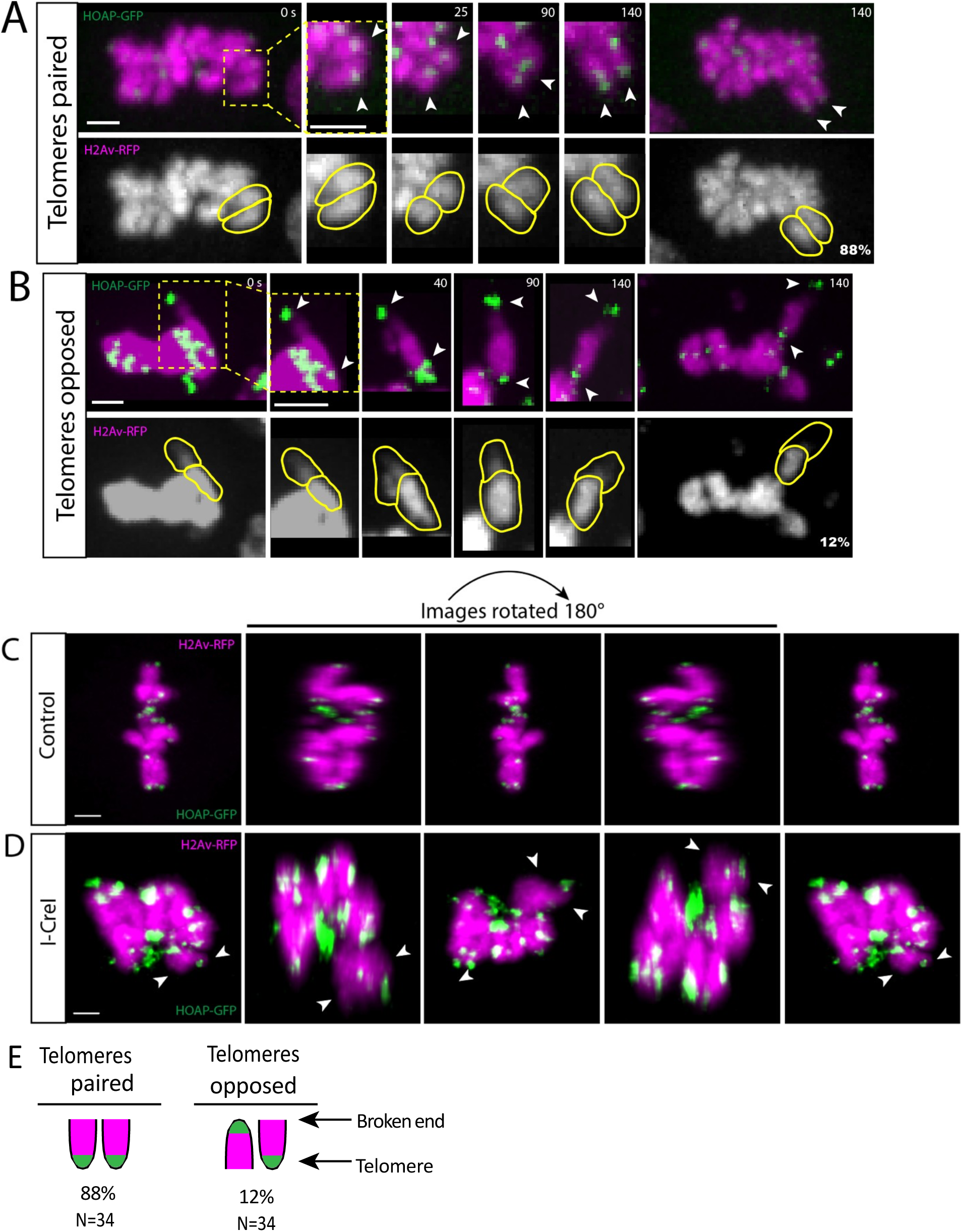
Acentrics align at the metaphase plate with telomeres either paired or unpaired. (A) Still frames of a time-lapse movie of a mitotic neuroblast labeled with H2AvRFP (magenta) and HOAP-GFP (green) expressing I-CreI with acentric sister telomeres aligned and paired (arrowheads and yellow outline) at metaphase (Video S5). (B) Still frames of a time-lapse movie of a mitotic neuroblast expressing I-CreI with acentric sister telomeres opposed (arrowheads and yellow outline) at metaphase. Most acentrics are oriented with their telomeres paired during metaphase. Bars, 2 μm. Time in seconds. (C) Still images of a 3D rendering of a mitotic neuroblast with H2Av-RFP (magenta) and HOAP-GFP (green) (N=5). (D) Still images of a 3D rendering of a mitotic neuroblast with I-CreI induced chromosome fragments (Video S6). All chromosomes are arranged in a semi-circular formation with distally oriented telomeres immediately prior to anaphase onset (N=10). Bars, 2 μm. Images are rotated 180°. (E) Schematic showing frequency of sister acentrics with telomeres paired vs. telomeres opposing.

### Acentric alignment requires microtubule plus-ends

Previous studies in *Drosophila* reported the microtubule-stabilizing protein Map205 and the microtubule plus-end associated protein EB1 are required for acentric sister separation during anaphase initiation [Vicars et al., 2021]. To determine if acentric alignment on the metaphase plate relies on microtubule plus-ends, we re-analyzed published data of cells expressing RNAi against the microtubule plus-end tracking protein EB1 [Vicars et al., 2021]. Our analysis reveals 27% (8/30) of acentric sisters unable to align with kinetochore-bearing chromosomes (Figure 5, Video S7). In control wild-type cells expressing only I-CreI, only 5% of acentric sisters fail to congress to the metaphase plate (Figure 5).

**Figure 5:**
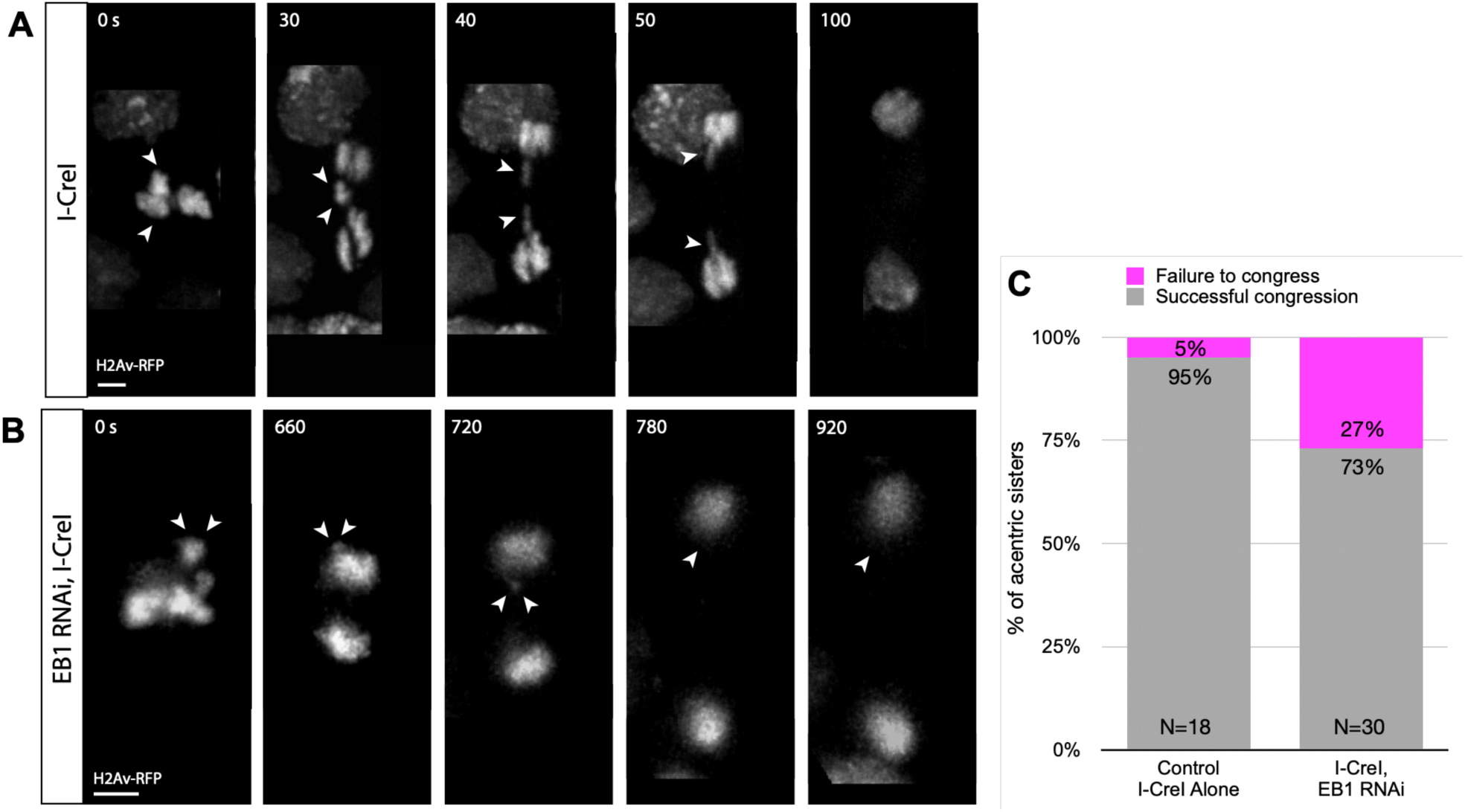
EB1 required for acentric metaphase plate alignment. (A) Still frames of a time-lapse movie of a mitotic neuroblast labeled with H2Av-RFP (gray) with I-CreI induced acentrics (white arrowheads). (B) Still frames of a time-lapse movie of a mitotic neuroblast expressing I-CreI and EB1 RNAi (Video S7). Sister acentrics fail to align with the chromosome mass at metaphase and segregate to the same cell pole. Bars, 2 μm. Time in seconds. (C) Bar graph showing congression failure rates. Successful chromosome congression is in gray. Failure of chromosomes to congress is in magenta.

A separate study demonstrated that peripheral interpolar microtubules play a critical role in the poleward segregation of acentric chromosome fragments [Karg et al., 2017]. Disruption of interpolar microtubule organization via knockdown of Klp3a (kinesin-4), disrupts the poleward segregation of the acentric, but not intact chromosomes. Additional laser-ablation experiments demonstrated that segregating acentrics are physically connected to the interpolar microtubules [Karg et al., 2017]. To determine if interpolar microtubules are involved in acentric movement to and alignment on the metaphase plate, we re-analyzed previously published acentric movies in a *klp3a* knockdown. *Drosophila* neuroblasts homozygous for hypomorphic alleles of *klp3a*, exhibited defects in congression of the acentrics, but not the intact chromosomes (Figure 6B & 6C, Video S8). In I-CreI expressing wild-type and *klp3a* mutant neuroblasts, 5% (1/18) and 30% (6/20) of acentrics fail to align with kinetochore-bearing chromosomes at the metaphase plate respectively (Figure 6C).

**Figure 6:**
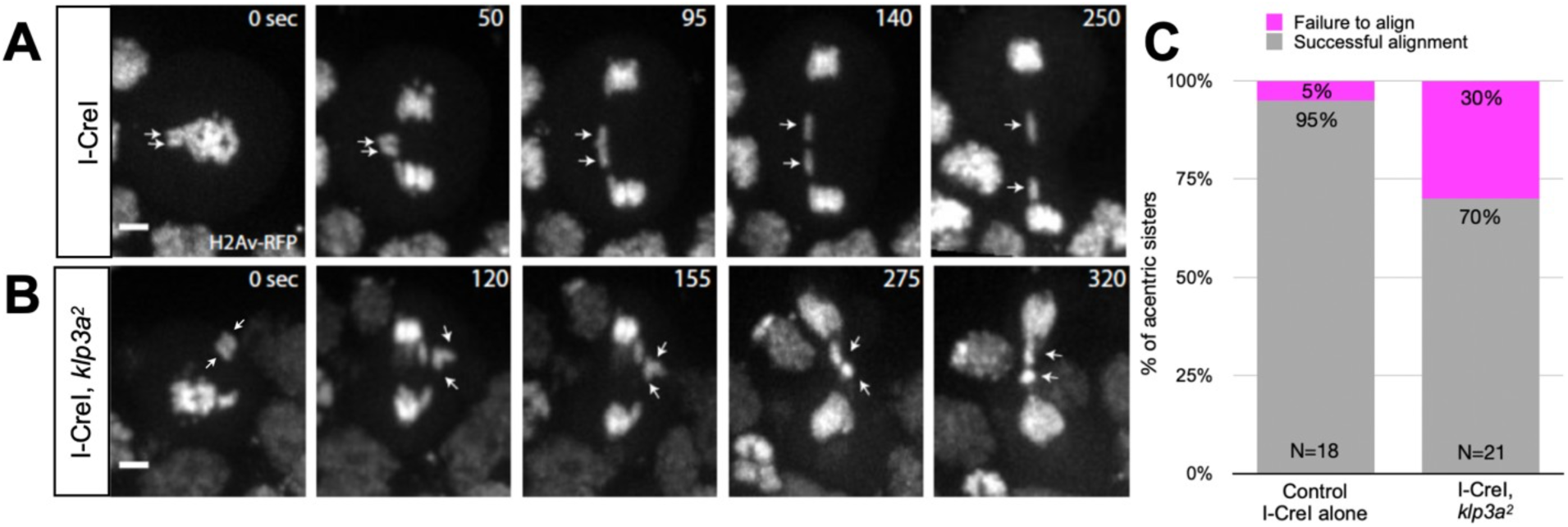
The chromokinesin Klp3a is required for aligning acentrics to the metaphase plate. (A) Still images from a time-lapse movie of a neuroblast expressing I-CreI alone (control). (B) Still images from a time-lapse movie of a neuroblast expressing I-CreI in *klp3a* mutant background (Video S8). Acentric sisters (white arrows) are positioned near the spindle pole at metaphase. Bars, 2 μm. Time in seconds. (C) Bar graph showing the rate of chromosome alignment at the metaphase plate. Successful chromosome alignment at the metaphase plate is in gray. Failure of chromosomes to align is in magenta.

Examination of chromosome dynamics on monopolar spindles provides a means to readily determine the extent to which microtubule plus-end forces act on chromosomes [Muscat et al., 2015]. Unfortunately, we discovered the small molecule inhibitors commonly used to induce monopolar spindles do not work in *Drosophila*. However, we find that RNAi knockdown of the chromokinesin Klp31E (kinesin 4) in *Drosophila* neuroblasts resulted in monopolar spindle formation. In these *Drosophila* neuroblasts with monopolar spindles, the acentric chromosomes, as well as the intact chromosomes, are pushed to the cell periphery (Figure 7). This finding indicates acentric chromosomes can be driven to microtubule plus ends prior to the metaphase-to-anaphase transition without requiring kinetochores. It remains to be determined if the acentrics are directly interacting with microtubule plus ends or utilizing motor proteins to travel to the microtubule plus ends.

**Figure 7:**
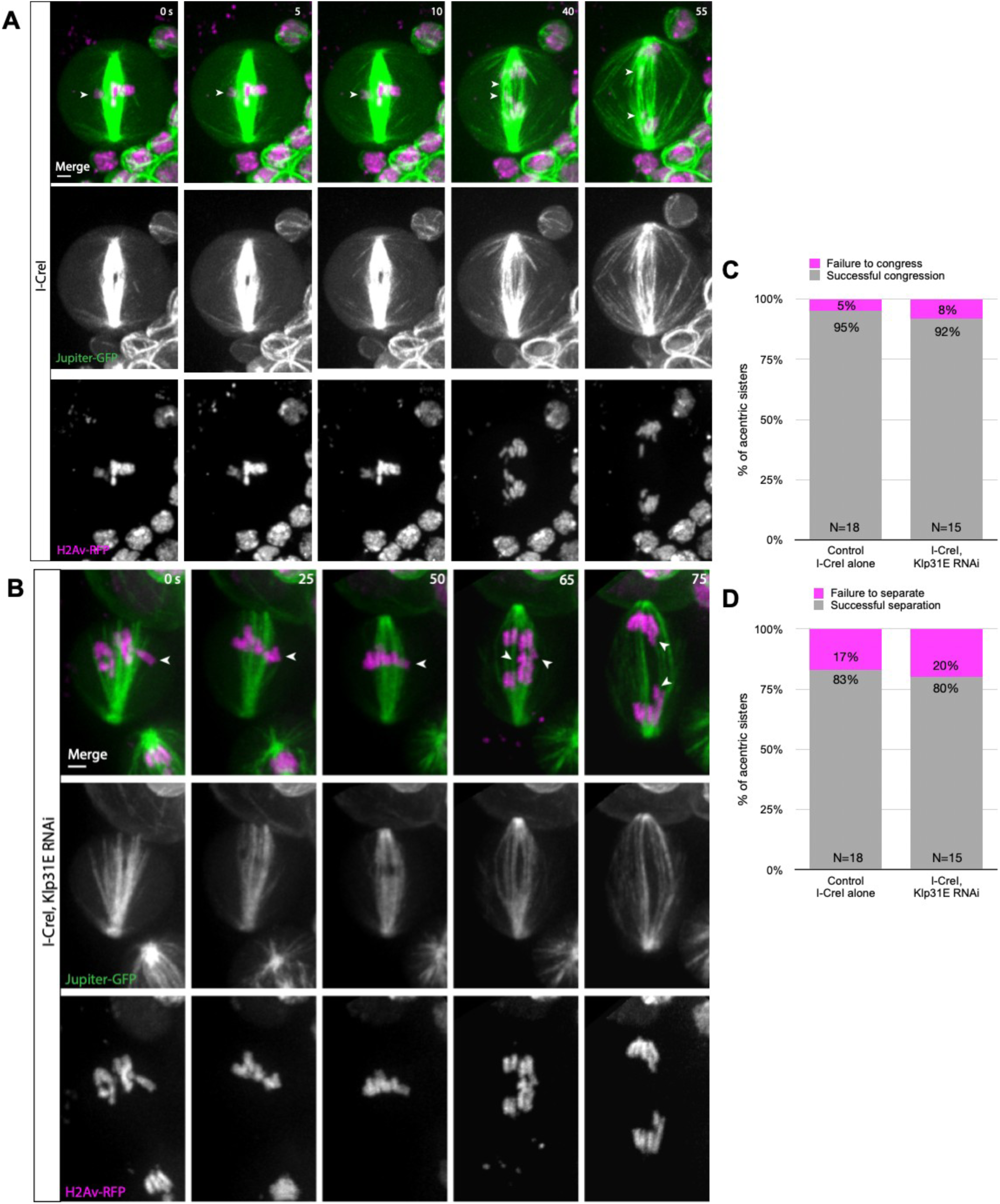
Acentrics are pushed by microtubule plus-ends. (A) Still frames of a time-lapse movie of a mitotic neuroblast expressing H2AvRFP (magenta) and Jupiter-GFP-labeled microtubules (green) with I-CreI induced acentrics. Sister acentrics (white arrowheads) align on the metaphase plate and lag behind at the spindle equator but eventually separate, segregate, and are incorporated into daughter nuclei. (B) Still frames of a time-lapse movie of a mitotic neuroblast with I-CreI induced acentrics (white arrowheads) and expressing Klp31E RNAi (Video S9). All chromosomes are pushed to the cell periphery by the mono-polar mitotic spindle. Bars, 2 μm. Time in seconds. (C) Percentages of acentric sisters that fail to congress to the metaphase plate. (D) Percentages of acentric sisters that fail to separate from one another during late anaphase.

**Figure 8:**
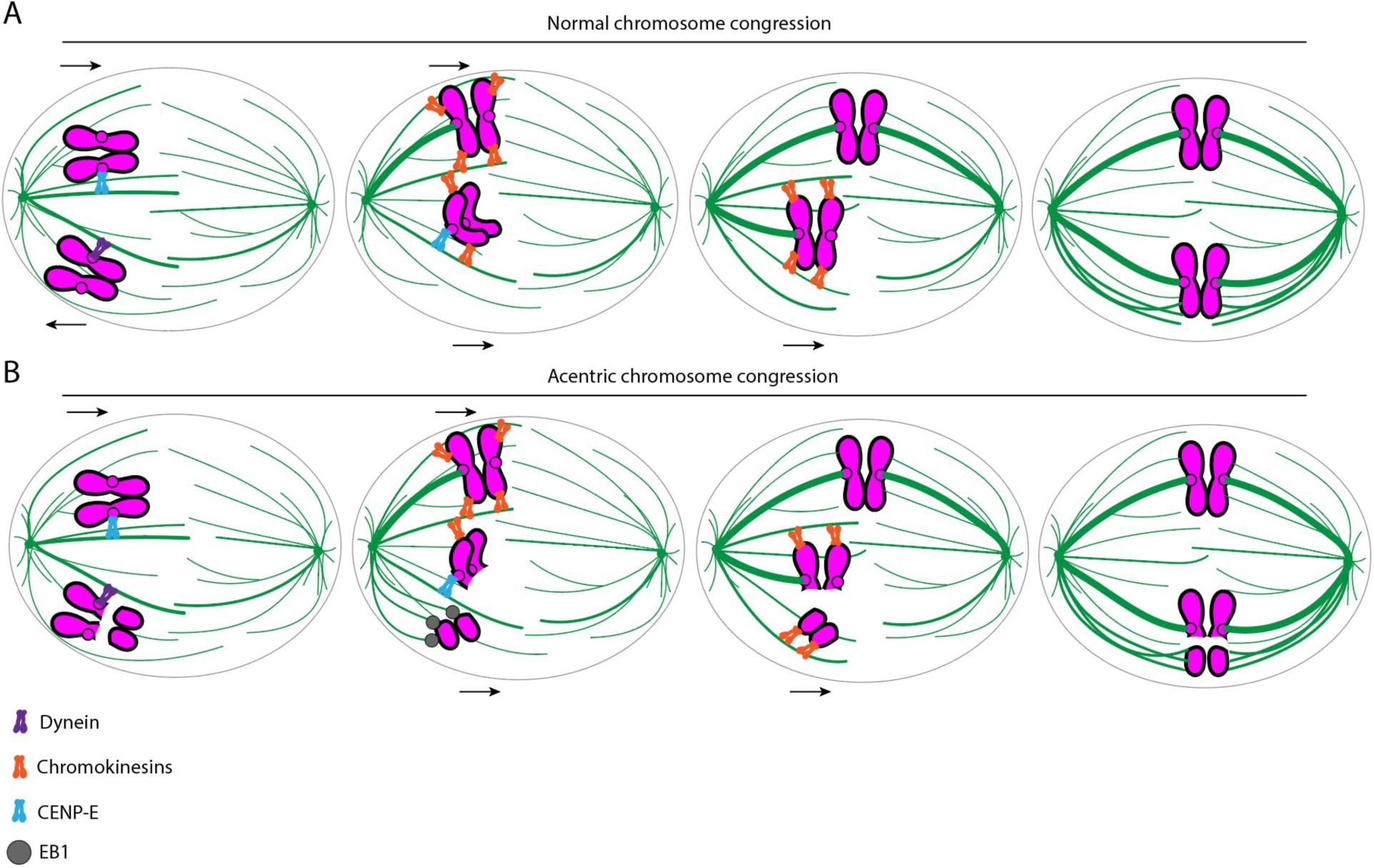
Schematic showing proposed mechanism of acentric chromosome congression. Chromosomes in magenta, microtubules in green, dynein in purple, chromokinesins in orange, CENP-E in blue, and EB1 in gray. (A) Normal chromosomes congress to the metaphase plate either through direct congression or peripheral congression. Direct congression relies on chromokinesins localized at chromosome arms and end-on kinetochore-microtubule aiachments. Peripheral congression involves chromosomes initially moving to the spindle pole by the minus-end motor protein dynein. Once at the pole, the chromosomes engage chromokinesins and the kinetochore-associated kinesin CENP-E for transport to the metaphase plate. (B) Acentric chromosome fragments congress by using EB1-driven microtubule plus-end pushing forces and lateral aiachments with microtubules (potentially via chromokinesins).

Surprisingly, despite the presence of monopolar spindles when Klp31E is partially knocked down, neuroblasts form an acentrosomal bipolar spindle allowing cells to successfully divide. 92% (14/15) of acentrics are able to align on the metaphase plate, with 80% (12/15) of acentric sisters successfully separating and segregating from one another and moving to opposite cell poles (Figure 7C & 7D).

## Discussion

Taking advantage of our ability to fluorescently tag the *Drosophila* X-chromosome acentric, we define the forces acting on the chromosome arms and the role of the kinetochore in chromosome congression. These studies revealed that during the late stages of congression, acentrics move faster than their neighboring kinetochore-bearing chromosomes. This indicates that kinetochore microtubule interactions act as a brake during congression slowing the action of plus-end directed forces acting on chromosome arms. In a similar fashion, the minus-end directed kinesin, Ncd (kinesin-14), acts as a brake on the action of the plus-end directed KLP61F (kinesin-5) in sliding anti-parallel microtubules during spindle pole separation [Tao et al., 2006]. Thus, the kinetochore serves to modulate and reduce the motion of the chromosomes during congression and alignment on the metaphase plate.

Our analysis also demonstrates that in contrast to intact chromosomes, the paired sister acentrics oscillate during the late stages of congression and upon arrival at the metaphase plate. The most likely explanation is that in the absence of a kinetochore, polar ejection forces unevenly distributed along the length of the chromosome generate oscillating movements. Thus, the kinetochore serves to stabilize chromosome orientation during alignment on the metaphase plate. As a result of these oscillations, sister acentrics orient either parallel or perpendicular to the spindle at anaphase onset. The former orientation results in acentric sister separation by sliding past one another while the later results in acentrics separating via “unzipping” with separation starting at one end of the chromosome [Vicars et al., 2021]

We also find that acentric sisters are positioned at the periphery of the metaphase plate and on a plane distinct from the rest of the chromosome complement. Previous studies demonstrated that the peripheral interpolar microtubules play a key role in acentric segregation during anaphase [Karg et al., 2017]. The positioning of the acentrics at the edge of the metaphase plate, and its association with interpolar microtubules, indicates that congression and alignment of the acentric chromosomes relies on interpolar microtubules. This conclusion is supported by the finding that knockdown of Klp3a, a kinesin required for establishing the population of interpolar microtubules, preferentially disrupts congression of acentric chromosomes [Karg et al., 2017]. *Drosophila* neuroblasts homozygous for hypomorphic alleles of *klp3a*, exhibit a dramatic increase in acentric congression defects (Figure 6). Since Klp3a is a chromokinesin, it is also possible that it interacts directly with acentrics propelling them along interpolar microtubules to the metaphase plate. Our finding that acentrics oscillate argues against this as direct lateral interactions between the acentric and microtubules would inhibit oscillation. In addition, the fact that the acentrics move faster than intact chromosomes suggests that they are not restricted by direct interactions with motor proteins. Acentrics may preferentially rely on interpolar microtubules due to the fact that kinetochore microtubules maintain intact chromosomes in a limited central region of the spindle. Lacking a kinetochore, acentrics are shunted to the interpolar microtubules.

Our studies also demonstrate that polar ejection forces play significant roles in acentric congression. This is most dramatically illustrated by the fact that reduction of EB1 activity preferentially disrupts alignment of the acentric on the metaphase plate but not the intact chromosomes. EB1 has been found to be essential for generating anti-polar forces on chromosomes and stabilizing microtubules [Tirnauer & Bierer, 2000; Vitre et al., 2008]. This finding is also supported by the finding that in monopolar spindles acentrics are rapidly pushed away from the poles. As described above, the oscillation of acentrics as they align at metaphase is readily explained by distributed plus-end directed forces acting along the chromosome arms.

One of the most surprising findings of this study emerges from live imaging paired sister acentrics with GFP-labeled telomeres. The labeled telomeres enable us to determine the relative orientation of each sister chromatid in the acentric pair as they align to the metaphase plate. As expected, intact sister chromatids exhibit parallel alignment across the entire length of the chromosome. This gene-for-gene alignment of paired sisters likely facilities their stable cohesion. In addition, it is essential for successful mitotic recombination [Lisby & Rothstein, 2015; Cohen-Fix, 2001]. Surprisingly, we find that a small but significant fraction of sister acentrics align in an anti-parallel orientation, indicating that the kinetochore is required to prevent antiparallel alignment of sister chromatids. The mechanism by which the kinetochore facilitates parallel alignment remains unknown as it is likely established during interphase immediately following completion of S-phase [Skibbens, 2008].

Another unexpected finding from this analysis is that the presence of the acentric chromosome fragment induces a global reorganization of congressed chromosomes. In wild-type *Drosophila* neuroblasts, fully congressed chromosomes are roughly organized as a column at metaphase. However, in the presence of the acentric chromosome fragment, the chromosomes arrange themselves in a ring-like configuration. With a lack of chromatin in the center of the ring the congressed chromosomes form a torus. Whether it is the presence of the acentric or damaged DNA that causes the global reorganization of the congressed chromosome remains unclear. Interestingly, this torus configuration of chromosomes is well-documented in human cells at prometaphase and metaphase [Magidson et al., 2011; Paulson et al., 2021]. 3D-reconstructions of human prometaphase chromosomes and metaphase chromosomes reveal their arrangement in a circular shape with kinetochores organized around the center and chromosome arms pushed towards the cell periphery [Magidson et al., 2011; Paulson et al., 2021; Renda et al., 2022]. It should be noted that human cancer cell lines often contain damaged DNA [Lewis & Golsteyn, 2016; Valencia-González et al., 2019]. As with acentric induction in *Drosophila*, the presence of DNA damage may induce the toroidal shape of the chromosome mass at metaphase.

Studies in other species reveal distinct mechanisms can result in congressed chromosomes forming a torus. For example, a similar torus configuration of congressed chromosomes has been observed in diatoms. In these species, interpolar microtubules gather in the center of the cell at metaphase, connecting spindle poles, and push chromosomes to either side of the interpolar microtubule bundle [Pickett-Heaps et al., 1975]. Additionally, in prometaphase newt pneumocytes, bundles of keratin filaments push chromosomes to the cell periphery [Hayden et al., 1990]. However, in *Drosophila* neuroblasts expressing I-CreI, the mitotic spindle is arranged with microtubules connecting to congressed chromosomes in the torus formation and an absence of microtubules in the center of the torus (Supplemental Figure 3). Whether the torus configuration has functional consequences or is simply an outcome of the mechanisms by which the spindle is formed is unknown. Taken together, these data support a model in which acentrics congress to and align on the metaphase plate via plus-end microtubule interactions stabilized by outer pole-to-pole microtubules and microtubule-associated proteins.

## Materials and methods

### Fly stocks

All stocks were raised on standard *Drosophila* media at room temperature (20–22°C) as previously described [Royou et al., 2008]. For generating acentrics, a transgenic fly line bearing the I-CreI endonuclease under heat-shock 70 promoter were kindly provided by Kent Golic at The University of Utah.

### Live analysis of acentric behavior in *Drosophila* third instar neuroblasts

As previously described, acentric chromosome fragments were induced by I-CreI expression (under heat shock 70 promoter) in 3rd instar larvae by a 1-hour 37°C heat shock followed by a 1-hour recovery period at room temperature [Royou et al., 2010]. The larval brains from third instar larvae were then dissected in PBS and placed on a slide with 20 μl of PBS. A coverslip was placed on the brain in PBS and the excess PBS was wicked out from the edges of the coverslip to induce squashing of brain between the slide and coverslip. Then, the edge of coverslip was sealed with halocarbon and was imaged for live analysis as described below. Neuroblast divisions in images for Figures 1-4, 6 and 7, Table 1, and Supplemental Figure 1 were from male 3rd instar larvae. Neuroblast divisions in images for Figure 5 and Supplemental Figure 2 were from female 3rd instar larvae.

### Microscopy and image acquisition

#### Wide-field microscopy

Time-lapse imaging for Figures 5B and 5C was performed using a Leica DM16000B wide-field inverted microscope equipped with a Hamamatsu electron-multiplying charge coupled device camera (ORCA 9100–02) with a binning of 1 and a 100x Plan-Apochromat objective with NA 1.4. Successive time points were filmed at 20 second intervals. RFP (585 nm) and GFP (508 nm) fluorophores were imaged. Samples were imaged in PBS and at room temperature (20–22°C). Widefield images were acquired with Leica Application Suite Advanced Fluorescence Software and 3D deconvolved using AutoQuant X2.2.0 software.

#### Spinning-disk microscopy

Images in Figures 1-4, 5A, 6, and 7, and Supplemental Figures 1 and 2 were acquired with an inverted Nikon Eclipse TE2000-E spinning disk (CSLI-X1) confocal microscope equipped with a Hamamatsu electron-multiplying charge coupled device camera (ImageEM X2) with a 100X 1.4 NA oil-immersion objective. Samples were imaged in PBS and at room temperature (20–22°C). Images were acquired with MicroManager 1.4 software. Time-lapse fluorescent images of neuroblasts divisions were done with 120 and 100 ms exposures for GFP and RFP respectively with 0.5 μm Z-steps. Time-lapse videos with both GFP and RFP were done every 5 seconds and time-lapse movies with RFP alone were done every 5 seconds. Figures were assembled in Adobe Illustrator. Selected stills (both experimental and control) were processed with ImageJ (http://rsb.info.nih.gov/ij/)).

## Measurements

In Figure 2, circularity measurements of chromosomes (H2Av-RFP) were done using the circularity function in ImageJ of the region outlined around acentrics and the main mass of chromosomes. For statistical analyses, two-sided Mann-Whitney U tests were used. Two-sided Mann-Whitney U tests were performed in R (R Core Team) and Prism Version 8 (GraphPad Software). 3D renderings in Figures 2 and 4 and Supplemental Figure 2 were created using Imaris software.

## Acknowledgements

We would like to thank Dr. Susan Strome, Dr. William Saxton, and Dr. John Tamkun at the University of California, Santa Cruz for use of their equipment and for providing reagents. We thank Dr. Benjamin Abrams for his advice and technical expertise regarding microscopy experiments. We thank Dr. Kent Golic at The University of Utah for providing us with the transgenic fly line bearing the I-CreI endonuclease.

Funding for these studies was provided by the National Institutes of Health grant NIGMS-1R35GM139595 awarded to W. Sullivan.

The authors declare no competing financial interests.

