## Supplemental Materials for "Acentric chromosome congression and alignment on the metaphase plate via kinetochore-independent forces in *Drosophila*"

### SUPPLEMENTAL FIGURES

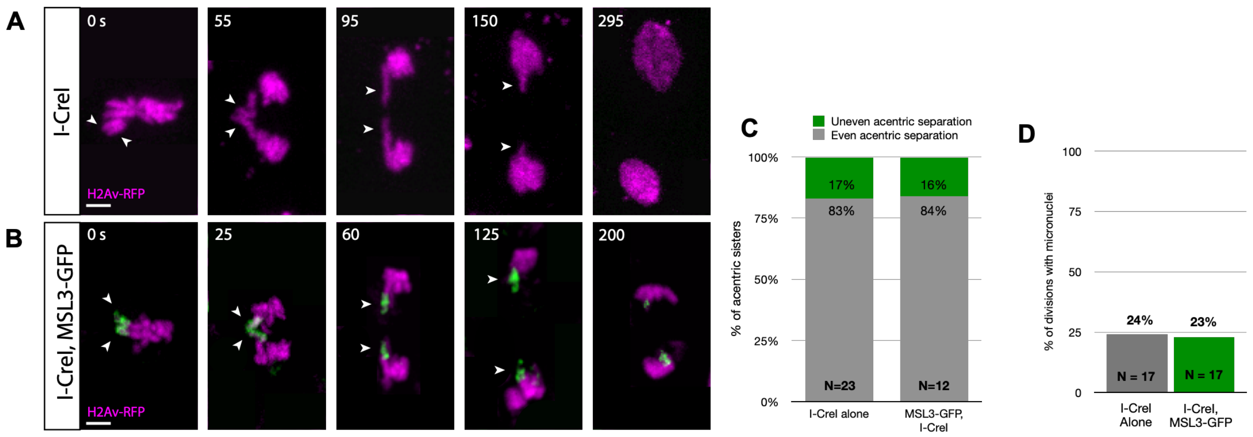


**Supplemental Figure 1: Neuroblasts expressing MSL3-GFP and I-CreI behave similarly to neuroblasts only expressing I-CreI.** (A) Still frames of a time-lapse movie of a mitotic neuroblast with I-CreI induced acentrics. Paired sister acentrics (white arrowheads) align at the metaphase plate, lag behind at the spindle equator in anaphase, and eventually separate, segregate, and incorporate into daughter nuclei. (B) Still frames of a time-lapse movie of a mitotic neuroblast with I-CreI induced acentrics and expressing MSL3-GFP. Bars, 2 μm. Time in seconds. (C) Percentages of acentric sisters that fail to completely separate from one another. (D) Percentages of neuroblast divisions in which acentrics failed to incorporate into daughter nuclei and formed one or more micronuclei.

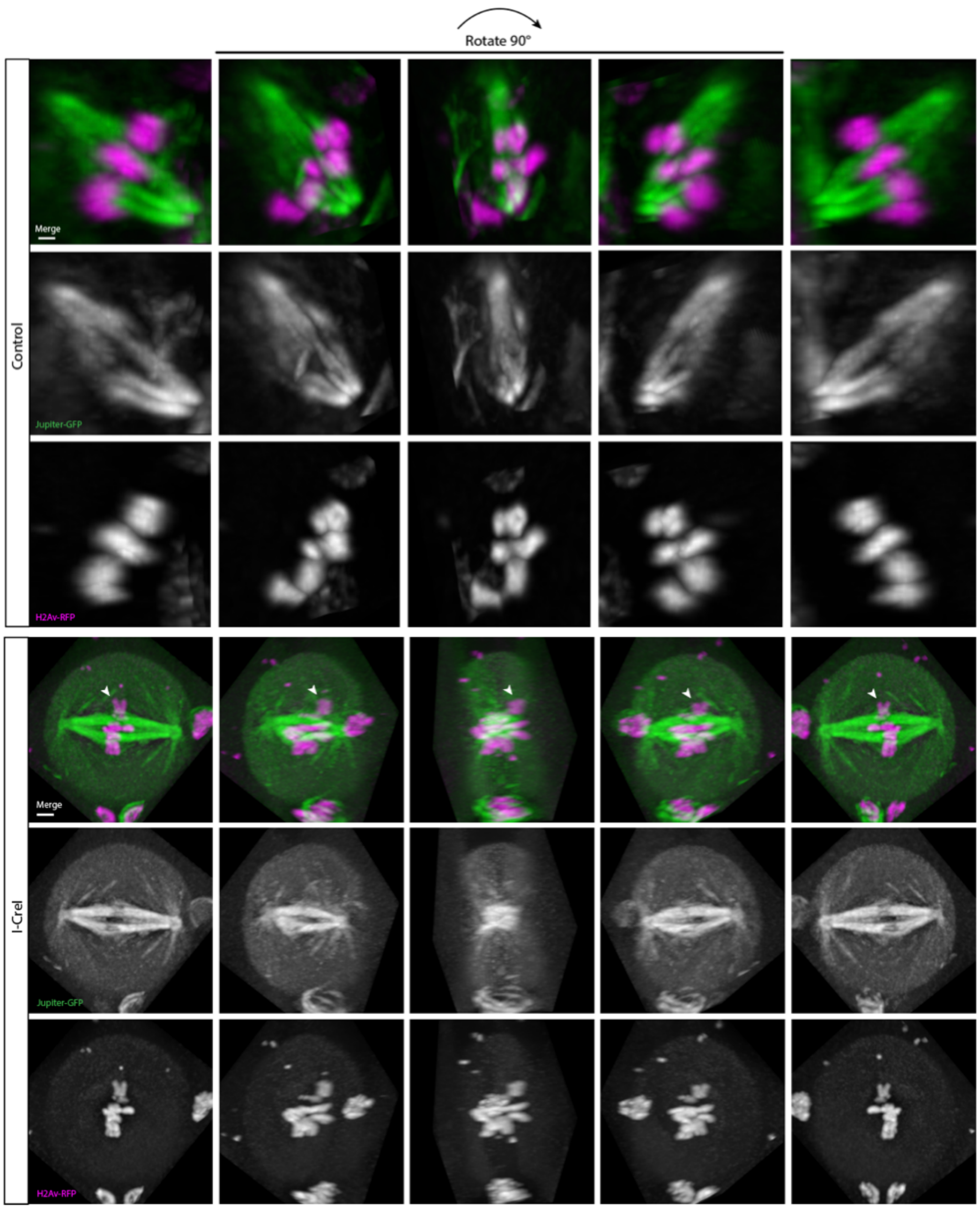


**Supplemental Figure 2: I-CreI induction produces a toroidal arrangement of congressed chromosomes with microtubules absent in the center.** A) Still images of a 3D rendering of a neuroblast at metaphase labeled with H2Av-RFP (magenta) and Jupiter-GFP labeled microtubules (green) not expressing I-CreI. (B) Still frames of a 3D rendering of a neuroblast at metaphase with I-CreI induced acentrics (white arrowheads). Bars, 2 µm. Images are rotated 90°.

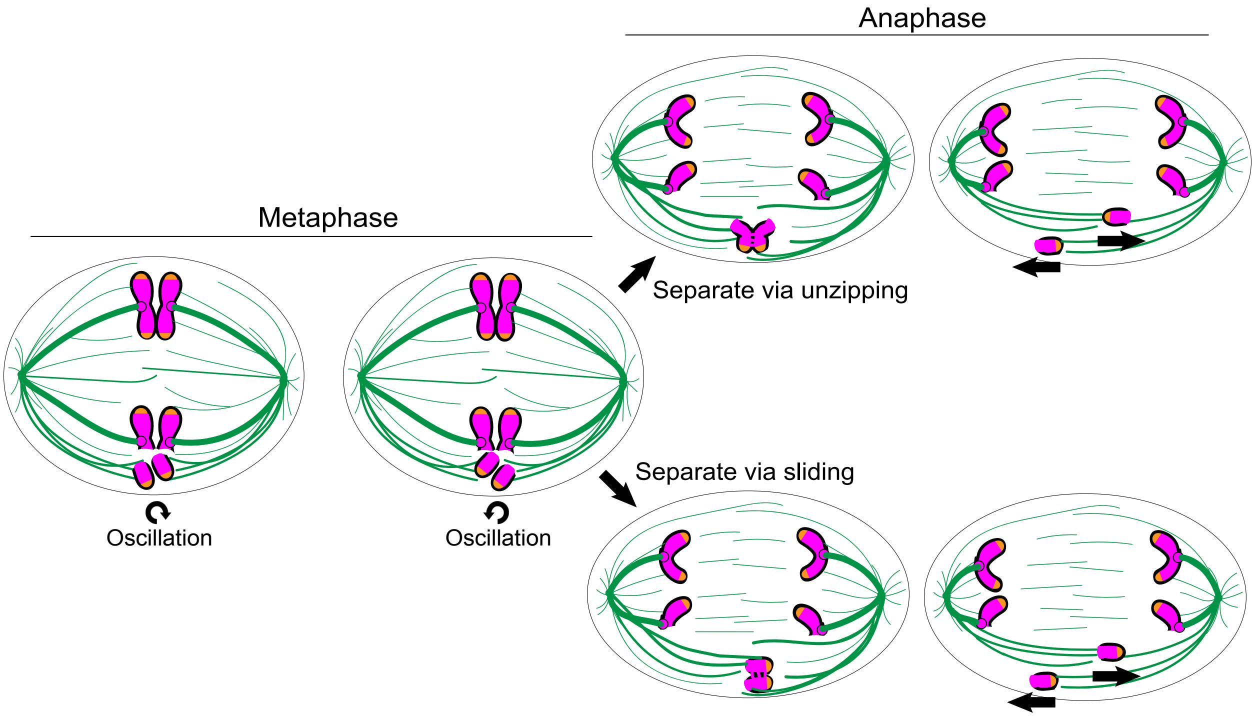


**Supplemental Figure 3: Schematic depicting acentric chromosome alignment at metaphase and separation in anaphase.** Chromosomes in magenta, microtubules in green, and telomeres in orange. Acentric sister chromatids align at the metaphase plate while undergoing oscillations. Depending on the final orientation of the acentrics upon anaphase onset, the acentrics may separate by “unzipping” from one another (perpendicular to the spindle) or by sliding past one another (parallel to the spindle). For further discussion of this model, see Vicars et al., 2021.
